# Meta-colored compacted de Bruijn graphs

**DOI:** 10.1101/2023.07.21.550101

**Authors:** Giulio Ermanno Pibiri, Jason Fan, Rob Patro

**Affiliations:** DAIS, Ca’ Foscari University of Venice, Venice, Italy; ISTI-CNR, Pisa, Italy; Department of Computer Science, University of Maryland, College Park, MD 20440, USA

## Abstract

**Motivation:** The colored compacted de Bruijn graph (c-dBG) has become a fundamental tool used across several areas of genomics and pangenomics. For example, it has been widely adopted by methods that perform read mapping or alignment, abundance estimation, and subsequent downstream analyses. These applications essentially regard the c-dBG as a map from *k*-mers to the set of references in which they appear. The c-dBG data structure should retrieve this set — the *color* of the *k*-mer — efficiently for any given *k*-mer, while using little memory. To aid retrieval, the colors are stored explicitly in the data structure and take considerable space for large reference collections, even when compressed. Reducing the space of the colors is therefore of utmost importance for large-scale sequence indexing.

**Results:** We describe the *meta-colored* compacted de Bruijn graph (Mac-dBG) — a new colored de Bruijn graph data structure where colors are represented holistically, i.e., taking into account their redundancy across the whole collection being indexed, rather than individually as atomic integer lists. This allows the factorization and compression of common sub-patterns across colors. While optimizing the space of our data structure is NP-hard, we propose a simple heuristic algorithm that yields practically good solutions. Results show that the Mac-dBG data structure improves substantially over the best previous space/time trade-off, by providing remarkably better compression effectiveness for the same (or better) query efficiency. This improved space/time trade-off is robust across different datasets and query workloads.

**Code availability:** A `C++17` implementation of the Mac-dBG is publicly available on GitHub at: https://github.com/jermp/fulgor.

## 1 Introduction

The colored compacted de Bruijn graph (c-dBG) has become a fundamental tool used across several areas of genomics and pangenomics. For example, it has been widely adopted by methods that perform read mapping or alignment, specifically with respect to RNA-seq and metagenomic identification and abundance estimation [1,2,3,4,5,6,7,8]; among methods that perform homology assessment and mapping of genomes [9,10]; for a variety of different tasks in pangenome analysis [11,12,13,14,15,16,17], and for storage and compression of genomic data [18]. In most of these applications, a key requirement of the underlying representation of the c-dBG is to be able to determine — with efficiency being critical — the set of references in which an individual *k*-mer appears. These motivations bring us to the following problem formulation.

### Problem 1 (Colored *k*-mer indexing)

Let ℛ = {*R*_1_, …, *R*_*N*_} be a collection of references. Each reference *R*_*i*_ is a string over the DNA alphabet *Σ* = {`A, C, G, T`}. We want to build a data structure (referred to as the *index* in the following) that allows us to retrieve the set Color(*x*) = {*i*|*x* ∈ *R*_*i*_} as efficiently as possible for any *k*-mer *x* ∈ *Σ*^*k*^. If the *k*-mer *x* does not occur in any reference, we say that Color(*x*) = ∅. Hereafter, we simply refer to the set Color(*x*) as the *color* of the *k*-mer *x*.

Of particular importance for biological analysis is the case where ℛ is a *pangenome*. Roughly speaking, a pangenome is a (large) set of genomes in a particular population, species or closely-related phylogenetic group. Pangenomes have revolutionized DNA analysis by providing a more comprehensive understanding of genetic diversity within a species [19,20]. Unlike traditional reference genomes, which represent a single individual or a small set of individuals, pangenomes incorporate genetic information from multiple individuals within a species or group. This approach is particularly valuable because it captures a wide range of genetic variations, including rare and unique sequences that may be absent from any particular reference genome.

### Contributions

The goal of this paper is to propose a solution to Problem 1 focusing on the specific, important, application scenario where ℛ is a pangenome. (We note, however, that the approaches described herein are general, and we expect them to work well on any corpus of highly-related genomes, whether or not they formally constitute a pangenome.) To best exploit the properties of Problem 1, we capitalize on recent indexing development for c-dBGs [21]. The result is the *meta-colored* compacted de Bruijn graph (Mac-dBG) — a new data structure where colors are represented *holistically*, i.e., taking into account their redundancy across the whole collection being indexed, rather than individually as atomic integer lists. After covering preliminary concepts in Section 2 and a review of the state of the art in Section 3, we describe the Mac-dBG in Section 4.1 and 4.2. We present the underlying NP-hard optimization problem in Section 4.3 and discuss a simple framework for constructing the Mac-dBG in Section 4.4. Section 5 presents experimental results to demonstrate that the Mac-dBG remarkably improves the best previous space/time trade-off in the literature. In fact, it essentially combines the space effectiveness of the most compact solutions with the query efficiency of the fastest solutions, at the expense of a slower construction algorithm. We conclude in Section 6 where we highlight some promising future directions.

A `C++17` implementation of the Mac-dBG is available at: https://github.com/jermp/fulgor.

## 2 Preliminaries: modular indexing of colored compacted de Bruijn graphs

In this section we provide some background information to better understand the design principles of the solutions we review in Section 3 and of the new one we propose in Section 4.

In principle, Problem 1 could be solved using a classic data structure from Information Retrieval — the *inverted index* [22,23]. In the context of this problem, the indexed documents are the references {*R*_1_, …, *R*_*N*_} in the collection ℛ and the terms of the inverted index are all distinct *k*-mers of ℛ. Using the notation from Problem 1, it follows that Color(*x*) is the inverted list of the term *x*. Let ℒ denote the inverted index for ℛ. The inverted index ℒ explicitly stores the ordered set Color(*x*) for each *k*-mer *x* ∈ ℛ. The goal is to implement the map *x* → Color(*x*) as efficiently as possible in terms of both memory usage and query time. To this end, all the distinct *k*-mers of ℛ are stored in an *associative* dictionary data structure 𝒟. Suppose we have *n* distinct *k*-mers in ℛ. These *k*-mers are stored losslessly in 𝒟. To implement the map *x* → Color(*x*), 𝒟 is required to support the operation Lookup(*x*), which returns ⊥ if *k*-mer *x* is not found in the dictionary or a unique integer identifier in [*n*] = {1, …, *n*} if *x* is found. Problem 1 can then be solved using these two data structures — 𝒟 and ℒ — thanks to the interplay between Lookup(*x*) and Color(*x*): logically, the index stores the sets {Color(*x*)}_*x*∈ℛ_ in some compressed form, *sorted by* the value of Lookup(*x*).

To exploit at best the potential of this modular decomposition into 𝒟 and ℒ, it is essential to rely on the specific properties of Problem 1. For example, we know that consecutive *k*-mers share (*k* − 1)-length overlaps; also, *k*-mers that co-occur in the same set of references have the same color. A useful, standard, formalism that captures these properties is the so-called *colored (compacted) de Bruijn graph* (c-dBG).

Let 𝒦 be the set of all the distinct *k*-mers of ℛ. The node-centric de Bruijn graph (dBG) of ℛ is a directed graph *G* (𝒦, *E*) whose nodes are the *k*-mers in 𝒦. There is an edge (*u, v*) ∈ *E* if the (*k* − 1)-length suffix of *u* equals the (*k* − 1)-length prefix of *v*. Note that the edge set *E* is implicitly defined by the set of nodes, and can therefore be omitted from subsequent definitions.

We refer to *k*-mers and nodes in a dBG interchangeably. Likewise, a path in a dBG spells the string obtained by concatenating together all the *k*-mers along the path, without repeating the shared (*k* − 1)-length overlaps. In particular, unary paths (i.e., non-branching) can be collapsed into single nodes spelling strings that are referred to as *unitigs*. Let 𝒰 = {*u*_1_, …, *u*_*m*_} be the set of unitigs of the graph. The dBG arising from this compaction step is called the *compacted* dBG, and indicated with *G* (𝒰).

The *colored* compacted dBG (c-dBG) is obtained by logically annotating each *k*-mer *x* with its color, Color(*x*). While different conventions have been adopted in the literature, here we assume that only non-branching paths with nodes having the *same* color are collapsed into unitigs. The unitigs of the c-dBG we consider in this work have the following key properties.

1. *Unitigs spell references in* ℛ. Each distinct *k*-mer of ℛ appears once, as sub-string of some unitig of the c-dBG. By construction, each reference *R*_*i*_ ∈ ℛ can be spelled out by some *tiling* of the unitigs — an ordered sequence of unitig occurrences that, when glued together (accounting for (*k*-1)-symbol overlap and orientation), spell *R*_*i*_ [24]. Joining together *k*-mers into unitigs reduces their storage requirements and accelerates looking up *k*-mers in consecutive order [25].
2. *Unitigs are monochromatic*. The *k*-mers belonging to the same unitig *u*_*i*_ all have the same color. We write *x* ∈ *u*_*i*_ to indicate that *k*-mer *x* is a sub-string of the unitig *u*_*i*_. Thus, we shall use Color(*u*_*i*_) to denote the color of each *k*-mer *x* ∈ *u*_*i*_.
3. *Unitigs co-occur*. Distinct unitigs often have the *same* color, i.e., they co-occur in the same set of references, because they derive from conserved sequences in indexed references that are longer than the unitigs themselves. We indicate with *z* the number of distinct colors 𝒞 = {*C*_1_, …, *C*_*z*_}. Note that *z* ≤ *m* and that, in practice, there are almost always *many more* unitigs than there are distinct colors.

Fig. 1a illustrates an example c-dBG with these properties. We refer to a compacted c-dBG as *G*(𝒰, 𝒞).

**Fig. 1:**
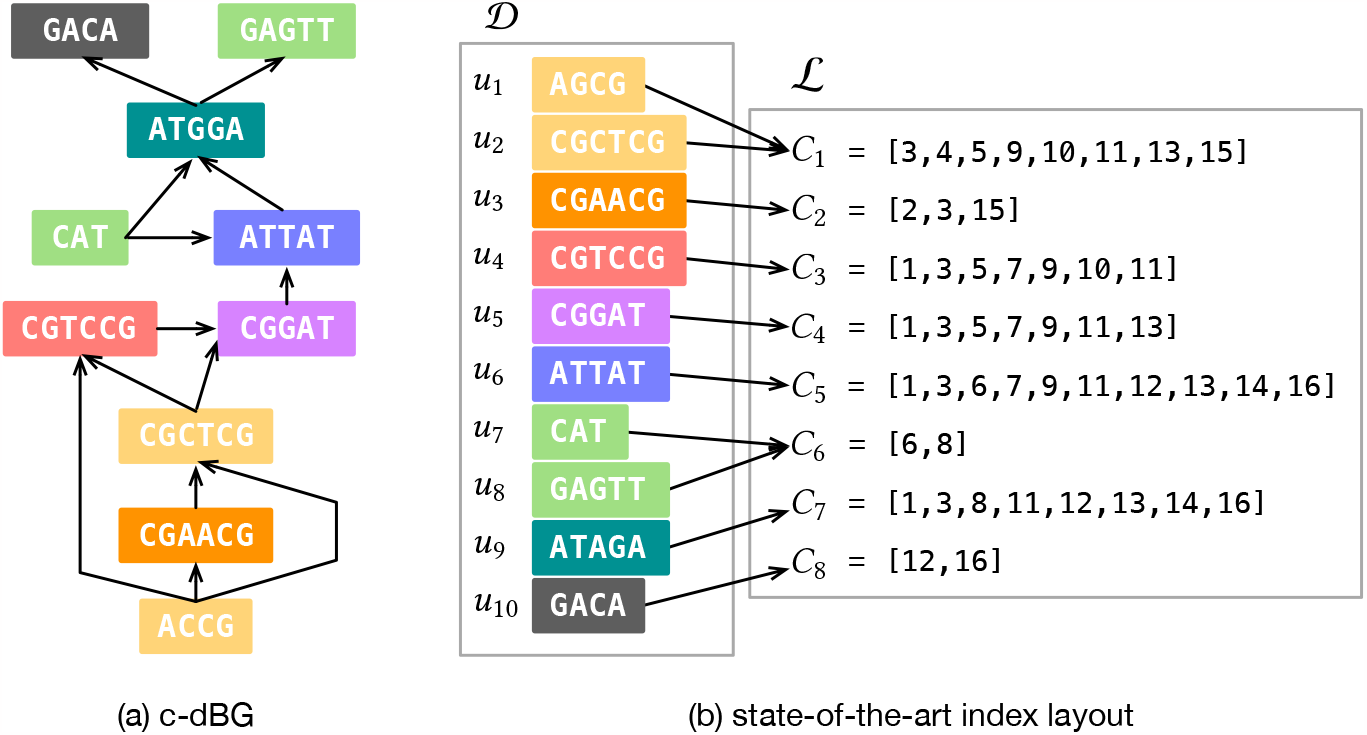
In panel (a), an example colored compacted de Bruijn graph (c-dBG) for *k* = 3. (In the figure, a *k*-mer and its reverse complement are considered as different *k*-mer for ease of illustration. In practice, these are considered identical.) The unitigs of the graph are colored according to the set of references they appear in. In panel (b), we schematically illustrate the state-of-the-art index layout (the Fulgor index [21]; see Section 3) assuming the c-dBG was built for *N* = 16 references, highlighting the modular composition of a *k*-mer dictionary, 𝒟, and an inverted index, ℒ. Note that unitigs are stored in 𝒟 in color order, hence allowing a very efficient mapping of *k*-mers to their distinct colors.

## 3 Related work

The solutions proposed in the literature to represent c-dBGs, and that fall under the “color-aggregative” classification [26], all provide different implementations of the modular indexing framework as described in Section 2. As such, they require an efficient *k*-mer dictionary along with a compressed inverted index.

For example, Themisto [27] makes use of the *spectral* BWT (SBWT) data structure [28] for its *k*-mer dictionary and an inverted index compressed with different strategies based on the sparseness of the color lists (ratio |*C*_*i*_| */N*). MetaGraph [29] uses the BOSS [30] data structure for the dictionary and exposes several general schemes to compress metadata associated with *k*-mers [29,31], which essentially constitute an inverted index. Bifrost [32], instead, uses a (dynamic) hash table to store the set of unitigs and an inverted index compressed with Roaring bitmaps [33]. The compact bit-sliced layout (or COBS) [34] can be considered as an *approximate* c-dBG in that the Color(x) query might contain false positives, i.e., spurious reference identifiers (but never false negatives). This is a consequence of building a Bloom filter for each reference, filled with all the *k*-mers in the reference. The Bloom filter matrix is stored in an inverted manner, and represents a collection of approximate colors. Being approximate, this method completely avoids the space consumption of the exact *k*-mer dictionary and the space is all due to the approximate colors.

However, none of these solutions simultaneously exploit all three unitig properties described in Section 2 to achieve faster query time and better space effectiveness. For example, Themisto disregards Property 1 as a direct consequence of using the SBWT data structure that internally arranges the *k*-mers in *colexicographic* order, and not in their order of appearance in the unitigs. This consideration is also valid of the BOSS data structure, hence for MetaGraph. Themisto exploits Property 3 instead, by compressing only the set of the *distinct* colors. Alanko et al. describe how it is possible in Themisto to reduce the space for the mapping from *k*-mers to colors by spending *O*(log *z*) bits for only some *k*-mers (the so-called *core k*-mers), while instead using 1 + *o*(1) bits for the other *k*-mers. However, this still requires dedicated storage related to the color *per* -*k*-mer, thus failing to exploit Property 2. Lastly, COBS does not exploit any specific property: unitigs are broken into their constituent *k*-mers and indexed separately; looking up consecutive *k*-mers (most likely part of the same unitig) has no locality of reference due to Bloom filter lookups; colors are stored approximately and partitioned into shards, so that a Color(x) query has to combine several partial results together.

To the best of our knowledge, the only solution that exploits *all* three properties is the recently-introduced Fulgor index [21], which we now review since it is the basis of our development in Section 4.

The solution implemented by Fulgor is to first map *k*-mers to unitigs using the dictionary 𝒟, and then succinctly map unitigs to their colors. The colors 𝒞 = {*C*_1_, …, *C*_*z*_} themselves are stored in compressed form in an inverted index ℒ. By *composing* these mappings, Fulgor obtains an efficient map directly from *k*-mers to their associated colors (see also Fig. 1b). The composition is made possible by leveraging the *order-preserving* property of its dictionary data structure — SSHash [25,35] — which explicitly stores the set of unitigs in *any* desired order. This property has some important implications. First, looking up consecutive *k*-mers is cache-efficient since unitigs are stored contiguously in memory as sequences of 2-bit characters. Second, if *k*-mer *x* occurs in unitig *u*_*i*_, the Lookup(*x*) operation of SSHash can efficiently determine the unitig identifier *i*, allowing to map *k*-mers to unitigs. Third, if unitigs are sorted in color order, so that unitigs having the same color are consecutive, then mapping a unitig to its color can be implemented in as little as 1 + *o*(1) bits *per unitig* and in constant time via a Rank query.

## 4 Meta-colored compacted de Bruijn graphs

When indexing large pangenomes, the space taken by the (compressed) colors dominates the whole index space [21,32,27] (see also the space breakdowns in the Supplementary material, Fig. 4a). Efforts toward improving the memory usage of c-dBGs should therefore be spent in devising better compression algorithms for the colors. In this work, we focus on exploiting the following crucial property that can enable substantially better compression effectiveness: *The genomes in a pangenome are very similar* which, in turn, implies that the *colors are also very similar* (albeit distinct).

By “similar” colors we mean that they share many (potentially, very long) identical integer sub-sequences. This property is not exploited if each color *C*_*i*_ is compressed *individually* from the other colors. For example, if *C*_*i*_ shares a long sub-sequence with *C*_*j*_, this sub-sequence is actually represented *twice* in the index, which wastes space. This example is instrumentally simple; yet, it suggests that the identification of such common sub-sequences across a large collection, as well as the design of an effective compression mechanism for these patterns, is not easy. A further complicating matter is that the example clearly generalizes to more than two sub-sequences, hence increasing with pangenome redundancy and aggravating the memory usage of an index that encodes them redundantly in each color.

To address this issue, we describe here the meta-colored compacted de Bruijn graph, or Mac-dBG. In the Mac-dBG, a color is represented as a sequence of references to sub-sequences that are shared with potentially many other colors. We refer to these references as *meta colors*. These common sub-sequences, which we call *partial colors*, are encoded once, rather than a number of times equal to the number of colors in which they appear. This allows reducing the required space for the index while incurring low query overhead when partial colors are sufficiently long. Indeed, we demonstrate experimentally in Section 5 that the Mac-dBG substantially improves over the space/time trade-off of a traditional c-dBG data structure.

Another key strength of this representation via meta/partial colors is its generality: it can applied to any c-dBG data structure arising from the composition of 𝒟 and ℒ to readily improve its space and query time.

### 4.1 Definition

Let *G*(𝒰, 𝒞) be the c-dBG built from the reference collection ℛ = {*R*_1_, …, *R*_*N*_}. We recall from Section 2 that we indicate with 𝒞 = {*C*_1_, …, *C*_*z*_} the set of distinct colors of *G*.

Let 𝒩 = {𝒩_1_, …, 𝒩_*r*_} be a partition of [*N*] = {1, …, *N*} for some *r* ≥ 1, i.e., 𝒩_*i*_ ≠ ∅ for all *i*, 𝒩_*i*_∩𝒩_*j*_ = ∅ for all (*i, j*) such that *i* ≠ *j*, and ∪𝒩_*i*_ = [*N*]. Let an order between the elements of each 𝒩_*i*_ = {*e*_*i,j*_} be fixed (for example, by sorting the elements in increasing order). Any 𝒩 induces a permutation *π* : [*N*] → [*N*], defined as *π*(*e*_*i,j*_) := *j* +*B*_*i*−1_ where 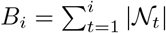 for *i >* 0 and *B*_0_ = 0, for *i* = 1, …, *r* and *j* = 1, …, |𝒩_*i*_|. We assume from now on that the *N* reference identifiers and the colors in 𝒞 have been permuted according to *π*. After the permutation, 𝒩 determines a partition of ℛ into *r* disjoint sets:

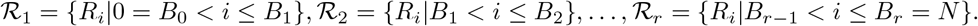

#### Definition 1 (Partial colors).

Let P_*i*_ be the set

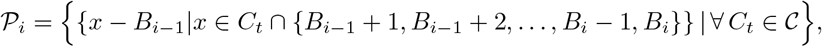

for *i* = 1, …, *r*. The elements {*P*_*ij*_} of the set 𝒫_*i*_ are the *partial colors* induced by the partition 𝒩_*i*_. We indicate with 𝒫 = {𝒫_1_, …, 𝒫_*r*_} the set of all partial color sets.

In words, 𝒫_*i*_ is the set obtained by considering the distinct colors *only* for the references in the *i*-th partition ℛ_*i*_ by noting that — by construction — they comprise integers *x* such that *B*_*i*−1_ *< x* ≤ *B*_*i*_.

The idea is that the set 𝒫 = {𝒫_1_, …, 𝒫_*r*_} form a dictionary of sub-sequences (the partial colors) that spell the original colors 𝒞 = {*C*_1_, …, *C*_*z*_}. Let us now formally define this spelling.

#### Definition 2 (Meta colors).

Let *C*_*t*_ ∈ 𝒞 be a color. A *meta color* is an integer pair (*i, j*) indicating the sub-list *L* := *C*_*t*_[*b* … *b* + |*P*_*ij*_|] if there exists 0 *< b* ≤ |*C*_*t*_| − |*P*_*ij*_| such that *L*[*l*] = *P*_*ij*_[*l*] + *B*_*i*−1_, for *l* = 1, …, |*P*_*ij*_|. It follows that *C*_*t*_ can be modeled as a list *M*_*t*_ of at most *r* meta colors. We indicate with M = {*M*_1_, …, *M*_*z*_} the set of all meta color lists.

Given *G*(𝒰, 𝒞), the Mac-dBG is the graph *G*(𝒰, 𝒩, *π*, 𝒫, ℳ) where the set of nodes, 𝒰, is the same as that of *G* but the colors 𝒞 are represented with the partial colors 𝒫 and the meta colors ℳ.

The Mac-dBG permits to encode the colors in 𝒞 into smaller space compared to the original c-dBG and without compromising the efficiency of the Color(*x*) query, for the following reasons.

1. If 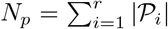 is the total number of partial color sets, then each meta color (*i, j*) can be indicated with just log_2_(*N*_*p*_) bits. Potentially long sub-lists, shared between several color lists, are therefore encoded once in 𝒫 and only referenced with log_2_(*N*_*p*_) bits instead of redundantly replicating their representation.
2. Each partial color *P*_*ij*_ can be encoded more succinctly because the permutation *π* guarantees that it only comprises integers lower-bounded by *B*_*i*−1_ + 1 and upper-bounded by *B*_*i*_. Hence only log_2_(*B*_*i*_ − *B*_*i*−1_) bits per integer are sufficient.
3. The total number of integers in 𝒫 is *less* than that in the original 𝒞, i.e., 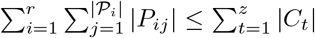 because partial colors are encoded once.
4. It is efficient to recover the original color *C*_*t*_ from the meta color list *M*_*t*_: for each meta color (*i, j*) ∈ *M*_*t*_, sum *B*_*i*−1_ back to each decoded integer of *P*_*ij*_. Hence, we decode *strictly increasing* integers. This is, again, a direct consequence of having permuted the reference identifiers with *π*. Observe that, in principle, the representation of the colors with meta/partial colors could be described *without* any permutation *π* — however, one would sacrifice space (for the reason 2. above) *and* query time since decoding a color list from meta colors would eventually need to sort the decoded integers. In conclusion, permuting the reference identifiers with *π* is an extra degree of freedom that we can exploit to improve index space and preserve query efficiency, noting that the correctness of the index is not compromised when reference identifiers are re-assigned globally.

#### Example 1.

Let us consider the *z* = 8 colors from Fig. 1b, for *N* = 16. Let *r* = 4 and 𝒩_1_ = {1, 12, 13, 14, 16}, 𝒩_2_ = {3, 5, 9}, 𝒩_3_ = {7, 11}, 𝒩_4_ = {2, 4, 6, 8, 10, 15}, assuming we use the natural order between the integers to determine an order between the elements of each 𝒩_*i*_. Thus, we have *B*_1_ = 5, *B*_2_ = 8, *B*_3_ = 10, and *B*_4_ = 16. The induced permutation *π* can be visualized by concatenating the sets _*i*_ from *i* = 1 to 4 and assigning “new” identifiers, from 1 to *N*, in this concatenated order:

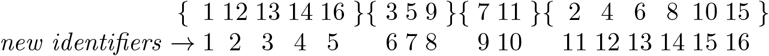

which results in *π*(1) = 1, *π*(12) = 2, *π*(13) = 3, etc., that is *π* = [1, 11, 6, 12, 7, 13, 9, 14, 8, 15, 10, 2, 3, 4, 16, 5]. Now we apply the permutation *π* to each color, obtaining the following permuted colors (vertical bars represent the partial color boundaries *B*_1_, …, *B*_4_).

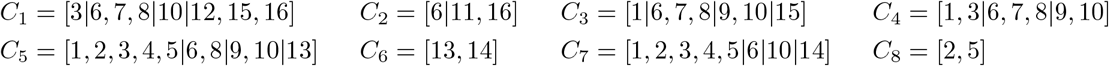

For example, color *C*_1_, that before was [3, 4, 5, 9, 10, 11, 13, 15] (see Fig. 1b), now is

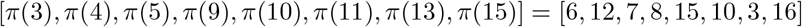

or [3, 6, 7, 8, 10, 12, 15, 16] once sorted. The partial colors are the distinct sub-sequences in each partition of the permuted colors. For example, 𝒫_1_ is the set of the distinct sub-sequences in partition 1, i.e., those comprising the integers *x* such that 0 *< x* ≤ *B*_1_ = 5. Hence, we have five distinct partial colors in partition 1, and these are [3], [1], [1, 3], [1, 2, 3, 4, 5], and [2, 5]. Importantly, note that from the integers in partial colors from partition *i >* 1 we can subtract the lower bound *B*_*i*−1_. For example, from the integers in the partial color [6, 7, 8] from *C*_1_ in partition 2 we can subtract *B*_1_ = 5, hence obtaining [1, 2, 3]. Overall, we thus obtain that 𝒫 comprises four partial color sets, as shown in Fig. 2. The figure also shows the rendering of the colors 𝒞 = {*C*_1_, …, *C*_8_} via meta color lists, i.e., how each each color can be spelled by a proper concatenation of partial colors.

**Fig. 2:**
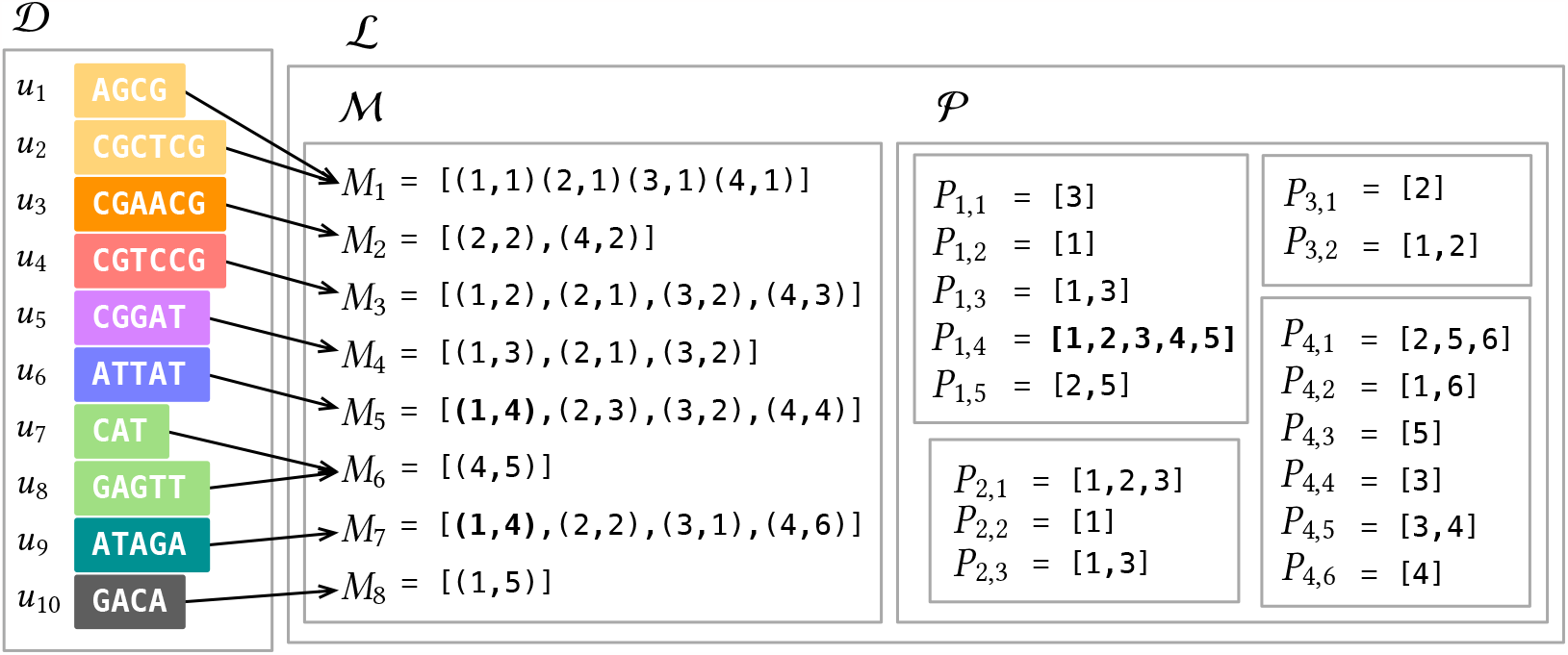
Mac-dBG layout discussed in Example 1 for the colors of the c-dBG from Fig. 1. Note that the partial color *P*_1,4_ = [1, 2, 3, 4, 5] shared between *C*_5_ and *C*_7_ is now represented *once* as a direct consequence of partitioning, and indicated with the pair (1, 4) instead of replicating the five integers it contains in both *C*_5_ and *C*_7_. The same consideration applies to other shared sub-sequences.

### 4.2 Data structures used and two-level intersection algorithm

Given a Mac-dBG *G*(𝒰, 𝒩, *π*, 𝒫, ℳ), a concrete implementation includes a representation for 𝒰, 𝒫, and ℳ (plus also the sorted array *B*[1..*r*] = [0, *B*_1_, …, *B*_*r*−1_]). The Mac-dBG is not bound to any specific compression scheme nor any specific dictionary data structure, allowing one to obtain a spectrum of different space/time trade-offs depending on choices made. In this paper, we made the following choices: (1) we use the SSHash data structure [25,35] to represent the set of unitigs 𝒰 ; (2) we adopt the same compression methods as used in Fulgor [21] to compress the partial colors and the same mechanism to map unitigs to their colors (using a binary vector of length *m*, equipped with ranking capabilities); (3) we represent each meta color list as a list of log_2_(*N*_*p*_)-bit integers.

Very importantly, note that choices (1) and (2) directly imply that our Mac-dBG implementation fully exploits the key unitig properties described in Section 2 as Fulgor does.

The Mac-dBG opens the possibility to achieve even faster query times than a traditional c-dBG, due to the manner in which the partitions factorize the space of references, if a two-level intersection algorithm is employed for pseudoalignment. There are several pseudoalignment algorithms (see [21, Section 4] for an overview) that standard c-dBG data structures directly support; here we focus on the *full intersection* algorithm. Given a query string *Q*, we consider it as a set of *k*-mers. Let 𝒦 (*Q*) = {*x* ∈ *Q*} Color(*x*) ≠ ∅. The full intersection method computes the intersection between the colors of all the *k*-mers in (*Q*). Our two-level intersection algorithm is as follows. First, only meta colors are intersected (thus, without any need to access the partial colors) to determine the partitions in common to all colors being intersected. Then only the common partitions are considered. Two cases can happen for each partition. (1) The meta color is the same for all colors: in this case, the result of the intersection is implicit and it suffices to decode the partial color indicated by the meta color. (2) The meta color is not the same, hence we have to compute the intersection between different partial colors. This optimization is beneficial when the colors being intersected have very few partitions in common, or when they have identical meta colors.

### 4.3 The optimization problem

As evident from its definition, the effectiveness of a Mac-dBG crucially depends on the choice of the partition 𝒩 and upon the order of the references within each partition as given by the permutation *π*. There i s, in fact, an evident friction between the encoding costs of the partial and meta colors. Let *N*_*m*_ and 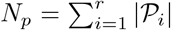 be the number of meta and partial colors, respectively. Since each meta color can be indicated with log_2_(*N*_*p*_) bits, meta colors cost *N*_*m*_ log_2_(*N*_*p*_) bits overall. Instead, let Cost(*P*_*ij*_, *π*) be the encoding cost (in bits) of the partial color *P*_*ij*_ according to some function Cost. On one hand, we would like to select a large value of *r* so that *N*_*p*_ diminishes since each color is partitioned into several, small, partial colors, thereby increasing the chances that each partition has many repeated sub-sequences. This will help in reducing the encoding cost for the partial colors, i.e., the quantity 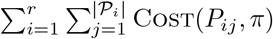. On the other hand, a large value of *r* will yield longer meta color lists, i.e., increase *N*_*m*_. This, in turn, could erode the benefit of encoding shared patterns and would require more time to decode each meta color list.

We can therefore formalize the following optimization problem that we call *minimum-cost partition arrangement* (MPA).

### Problem 2 (Minimum-cost partition arrangement)

Let *G*(𝒰, 𝒞) be the compacted c-dBG built from the reference collection ℛ = {*R*_1_, …, *R*_*N*_}. Determine the partition 𝒩 = {𝒩_1_, …, 𝒩_*r*_} of [*N*] = {1, …, *N*} for some *r* ≥ 1 and permutation *π* : [*N*] → [*N*] such that 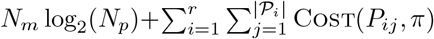 is minimum.

Depending upon the encoding scheme we choose, smaller values of Cost(*P*_*ij*_, *π*) may be obtained when the gaps between subsequent reference identifiers are minimized. Finding the permutation *π* that minimizes the gaps between the identifiers over all partial colors is an instance of the bipartite minimum logarithmic arrangement problem (BIMLOGA) as introduced by Dhulipala et al. [36] for the purpose of minimizing the cost of delta-encoded lists in inverted indexes. The BIMLOGA problem generalizes other known optimization problems, such as minimum logarithmic arrangement (MLOGA) and minimum logarithmic gap arrangement (MLOGGAPA) [37] problems, which are themselves modifications of the classic minimum linear arrangement problem (MLA) — all these problems are known to be NP-hard [38,39,37,36].

We note that BIMLOGA is a special case of MPA: that for *r* = 1 (one partition only) and Cost(*P*_*ij*_, *π*) being the log_2_ of the gaps between consecutive integers. It follows that also MPA is NP-hard under these constraints. This result immediately suggests that it is unlikely that polynomial-time algorithms exist for solving the MPA problem.

### 4.4 The SCPO framework

In this section we propose a construction algorithm for the Mac-dBG. The algorithm is an heuristic for the MPA optimization problem defined in the previous section (Problem 2), and it is based on the intuition that *similar* references should be grouped together in the same partition so as to *increase the likeliness of having a smaller number of longer shared sub-sequences*. The algorithm therefore consists in the following four steps: (1) *Sketching*, (2) *Clustering*, (3) *Partitioning*, and (4) *Ordering* (SCPO).

#### 1 Sketching

We argue that a reasonable way of assessing the similarity between two references is determining the number of unitigs that they have in common. Recall from Property 1 (Section 2) that each reference *R*_*i*_ ∈ ℛ can be spelled by a proper concatenation (a “tiling”) of the unitigs of the underlying compacted dBG. If these unitigs are assigned unique identifiers by SSHash, it follows that each *R*_*i*_ can be seen as a list of unitig identifiers. The idea is that these integer lists are much shorter and take less space than the actual DNA references. To reduce the space of a list even further, we compute a *sketch* of the list based on the fact that if two sketches are similar, then the original lists are similar as well.

#### 2 Clustering

The sketches are fed as input of a clustering algorithm.

#### 3 Partitioning

Once the clustering is done, each input reference *R*_*i*_ is labeled with the cluster label of the corresponding sketch so that the partition of ℛ into ℛ_1_, …, ℛ_*r*_ is uniquely determined.

#### 4 Ordering

Finally, one may *order* the references in each ℛ_*i*_ to determine a permutation *π* that yields a better compression for the partial colors 𝒫_*i*_. In fact, while the goal of clustering and partitioning is to factor out repeated sub-patterns within the colors, the goal of the ordering step is to assign nearby identifiers to references that tend to co-occur within the partial colors (as already mentioned in Section 4.3).

In this work, we use the following specific instance of this framework. We build *hyper-log-log* [40] sketches of *W* = 2^10^ bytes each. As clustering algorithm, we use a *divisive K*-means approach that does not need an a-priori number of clusters to be supplied as input. At the beginning of the algorithm, the whole input forms a single cluster that is recursively split into two clusters until the mean squared error (MSE) between the sketches in the cluster and the cluster’s centroid is not below a prescribed threshold (which we fix to 10% of the MSE at the start of the algorithm). Let *r* be the number of found clusters. The complexity of the algorithm depends on the topology of the binary tree representing the hierarchy of splits performed. In the worst case, the topology is completely unbalanced and the complexity is *O*(*WNr*); in the best case, the topology is perfectly balanced instead, for a cost of *O*(*WN* log *r*). Note that the worst-case bound is very pessimistic because, in practice, the formed clusters tend to be reasonably well-balanced in size.

In the current version of the work, we did not perform any ordering of the references within each cluster. We leave the investigation of this opportunity as future work.

Lastly in this section, it is worth noting that the approach we describe here for constructing Mac-dBGs bears a conceptual resemblance to the phylogenetic compression framework recently introduced by Břinda et al. [41]. At a high level, this owes to the fact that both approaches take advantage of well-known concepts in compression and Information Retrieval — namely that clustering and reordering are practical and effective heuristics for boosting compression. However, while the approach by Břinda et al. focuses on clustering references so as to improve the construction of collections of disparate dictionaries, we strictly focus on the effectiveness and efficiency of the index. As such, our approach adopts a single *k*-mer dictionary and instead induces a logical partitioning over the colors. This layout allows to avoid having to record *k*-mers that appear in multiple partitions more than once. As a result, while the phylogenetic compression framework aims to scale to immense and highly-diverse collections of references, it anticipates a primarily disk-based index in which partitions are loaded, decompressed, and searched for matches, similarly to a database search (or similarly to BLAST [42]). On the other hand, the Mac-dBG approach we present here places a premium on query time, and aims to enable *in-memory indexing* with interactive lookups for the purpose of fast read-mapping against the index.

## 5 Experiments

In this section we present the results of experiments conducted to assess the performance of the Mac-dBG. (The interested reader can find further experiments and details in the Supplementary material.) We fixed the *k*-mer length to *k* = 31. All experiments were run on a machine equipped with Intel Xeon Platinum 8276L CPUs (clocked at 2.20GHz), 500 GB of RAM, and running Linux 4.15.0.

### Datasets

We build Mac-dBGs with the proposed SCPO framework on the following pangenomes: 3,682 *E. Coli* (`EC`) genomes from NCBI [43]; different collections of *S. Enterica* (`SE`) genomes (from 5,000 up to 150,000 genomes) from the collection by Blackwell et al. [44]. Additionally, we also include a much more diverse collection of 30,691 genomes assembled from human gut samples (`GB`), originally published by Hiseni et al. [45]. Table 4 in the Supplementary material reports some basic statistics about these collections.

### Other evaluated tools

We compare the Mac-dBG against the following indexes, reviewed in Section 3: Fulgor [21], Themisto [27], MetaGraph [46,29,31], and COBS [34]. Links to the corresponding software libraries can be found in the References We use the C++ implementations from the respective authors. All software was compiled with gcc 11.1.0. We provide some details on the tested tools.

Both Themisto and COBS were built under default parameters as suggested by the authors, that is: option -d 20 for Themisto which enables the sampling of *k*-mer colors in the SBWT for better space effectiveness; in COBS, we have shards of at most 1024 references where each Bloom filter has a false positive rate of 0.3 and one hash function. MetaGraph indexes were built with the *relaxed row-diff* BRWT data structure [29] using a workflow available at https://github.com/theJasonFan/metagraph-workflows that we wrote with the input of the MetaGraph authors.

### Index size

Table 1 reports the total *on disk* index size for all of the methods evaluated. Compared to the most recent indexes, Fulgor and Themisto, that where previously shown to achieve the most desirable space/time trade-offs, Mac-dBG substantially improves on the space (and, as we shall see next, without any negative impact on query time). In fact, the only index smaller on disk than Mac-dBG is MetaGraph in the relaxed row-diff BRWT configuration — at least in the cases where we were able to construct the latter within the construction resource constraints. However, unlike the other indexes evaluated, the *on disk* index size MetaGraph is not representative of the working memory required for query when using the (recommended and default) batch mode query.

**Table 1:**
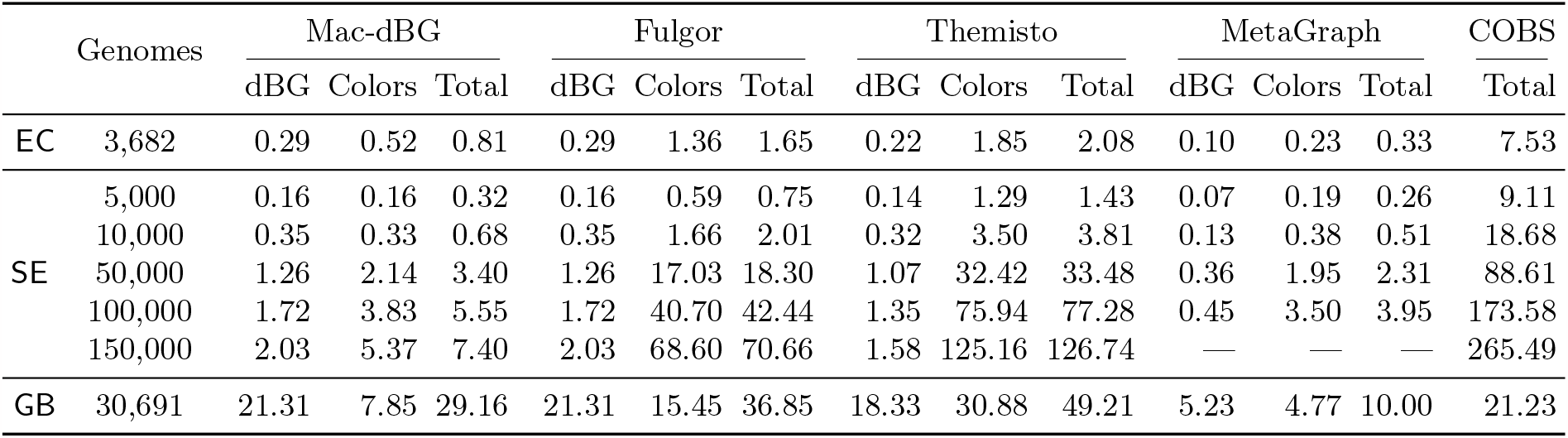
Index space in GB, broken down by space required for indexing the *k*-mers in the dBG (SSHash for both Fulgor and Mac-dBG, SBWT for Themisto, and BOSS for MetaGraph) and data structures required to encode colors and map *k*-mers to colors.

The COBS index, despite being approximate, is consistently and considerably larger than all of the other (exact) indexes, except for the the Gut bacteria collection (`GB`). The differing behavior on `GB` likely derives from the fact that the diversity of that data cause the exact indexes to spend a considerable fraction of their total size on the representation of the *k*-mer dictionary itself (e.g., 18 − 21.3 GB). However COBS, by design, eliminates this component of the index entirely.

Finally we observe that, as the number of references grow in the `SE` datasets, the already-large savings of Mac-dBG become even more prominent. For example Mac-dBG is 43% of the size of Fulgor (2.34× smaller) for `SE` 5,000, but is only 10% of the size of Fulgor (9.55× smaller) for `SE` 150,000. As the size of the collection grows, and more repetitive sub-patterns in the collection of colors appears, the Mac-dBG index is able to better capture and eliminate this redundancy.

### Query efficiency

Table 2 reports the query times of the indexes, performing full-intersection pseudoalignment (see Alg. 1 from [21]), on a high-hit workload. The performance on low-hit workloads is less informative, but is provided in the Supplementary material (Table 5) for completeness. The queried reads consist of all FASTQ records in the first read file of the following accessions: `SRR1928200` for `EC`, `SRR801268` for `SE`, and `ERR321482` for `GB`. These files contain several million reads each. Timings are relative to a second run of each experiment, where the indexes are loaded from the disk cache (which benefits the larger indexes more than the smaller ones).

**Table 2:**
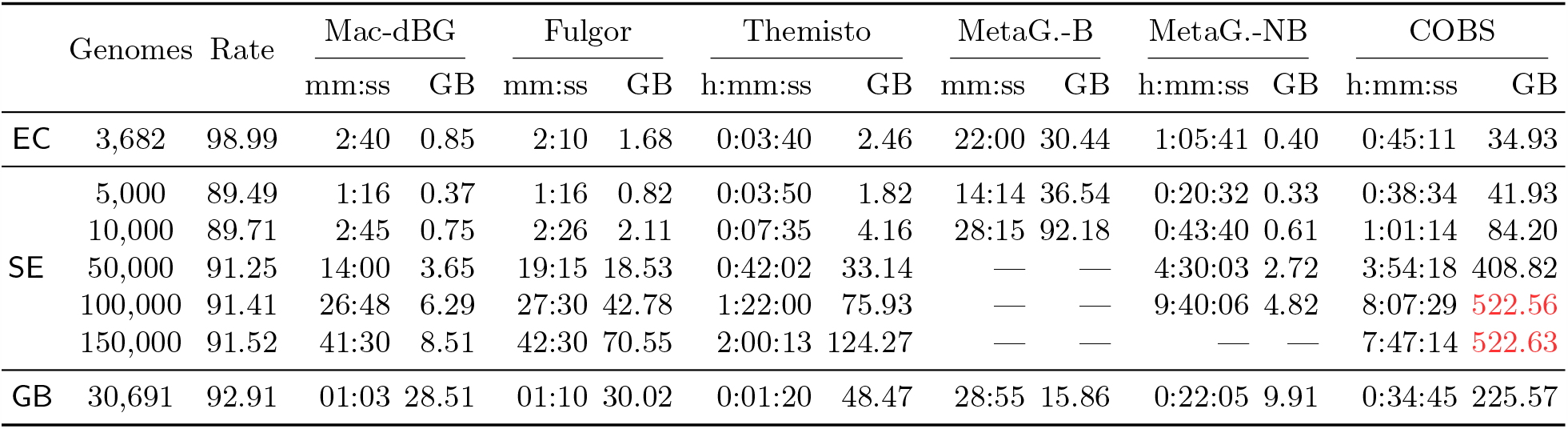
Total query time (elapsed time) and memory used during query (max. RSS) as reported by `/usr/bin/time -v`, using 16 processing threads. The read-mapping output is written to `/dev/null` for this experiment. We also report the mapping rate in percentage (fraction of mapped read over the total number of queried reads). The query algorithm used here is full-intersection. The “B” query mode of Meta-Graph corresponds to the batch mode (with default batch size); the “NB” corresponds to the non-batch query mode instead. In red font we highlight the workloads exceeding the available memory (*>* 500 GB).

Consistent with previously reported results [21], we find that among existing indexes, Fulgor provides the fastest queries. As expected, Mac-dBG does not not sacrifice query efficiency compared to Fulgor. After Mac-dBG and Fulgor, we note that Themisto is the next fastest index, followed by MetaGraph in batch query mode. The query speeds of COBS and of MetaGraph when not executed in batch mode are much lower than that of the other indexes, in some cases being (more than) an order of magnitude slower.

Critically, it is not the case with all indexes evaluated here that the size of the index *on disk* is a good proxy for the memory required to actually query the index. Specifically, for MetaGraph, when used in batch query mode (“B”), the required memory can exceed the on-disk index size by up to 2 orders of magnitude, and in several tests this resulted in the exhaustion of available memory and an inability to complete the queries under the tested configuration. On the other hand, Fulgor, Themisto, Mac-dBG and MetaGraph when not executed in batch mode (“NB”) require only a small constant amount of working memory beyond the size of the index present on disk.

### Construction time and space

In Table 3 we consider the resources needed to build the indexes. The Mac-dBG is built from a Fulgor index to which we apply the SCPO framework. For this reason, the time reported in Table 3 for the Mac-dBG has to be summed to the time needed to first build a Fulgor index. We do not describe here the construction algorithm for the Mac-dBG for space constraints; we just point out that the algorithm is single-threaded except for the construction of the sketches. Despite not being heavily engineered yet, the end-to-end construction of the Mac-dBG is competitive to that of Themisto and much faster than that of MetaGraph. As for the memory used, the end-to-end construction of the Mac-dBG takes the max. RSS between that of SCPO and that of Fulgor, which is always the max. RSS of Fulgor.

**Table 3:**
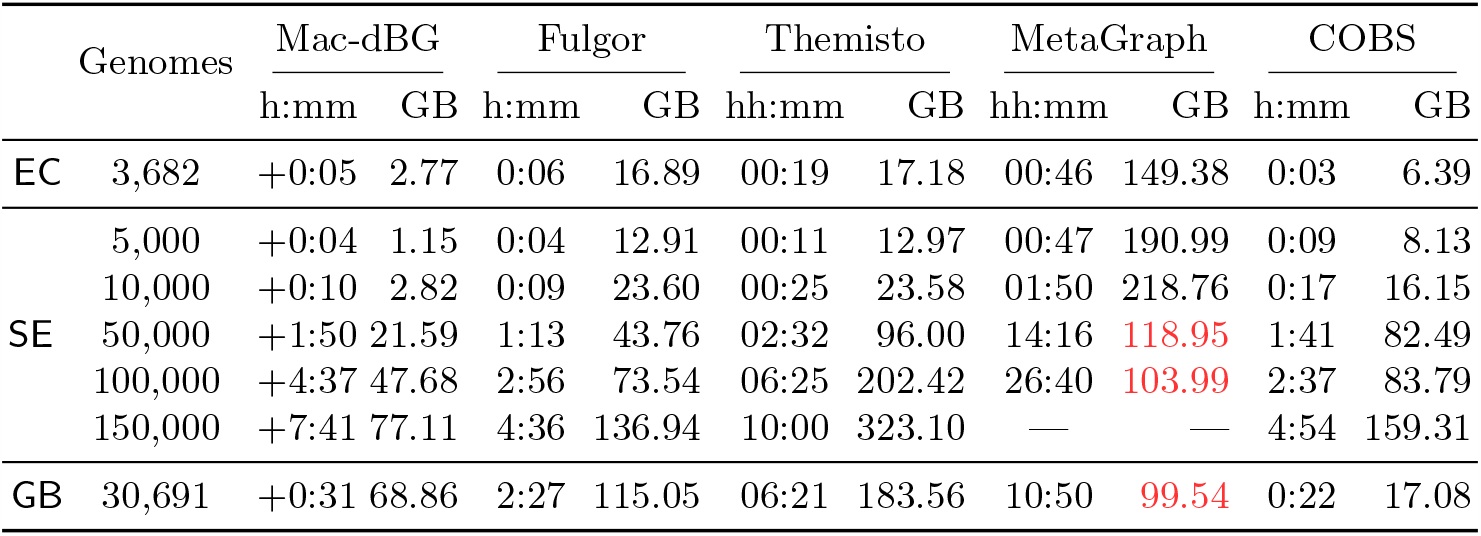
Total index construction time (elapsed time) and GB of memory used during construction (max. RSS), as reported by `/usr/bin/time -v`, using 48 processing threads. The reported time includes the time to serialize the index on disk. In red font we highlight the constructions exceeding the available memory (*>* 500 GB) and for which we had to cap the maximum memory usage to 100 GB. The time reported for the Mac-dBG is that for running the SCPO construction, hence it has to be summed to the time needed to first build a Fulgor index to partition.

The fastest indexes to build are Fulgor and COBS, the latter being even faster on the `GB` collection for reasons already explained. The tested MetaGraph configuration is significantly slower to build than all the other indexes; for example, we were unable to build the index for `SE` 150,000 within 3 days and using 48 parallel threads (the construction process also produced *>* 1 TB of intermediate files).

## 6 Conclusions

We have introduced the Mac-dBG data structure. The Mac-dBG represents a new state-of-the-art representation for answering color queries over large collections of reference sequences, and achieves a considerable improvement over existing work in terms of the space/time trade-off it offers. Specifically, Mac-dBG is almost as small as the smallest variant of MetaGraph — which is the smallest compressed c-dBG representation on disk. Yet, when queried, Mac-dBG requires essentially the same space as is required for the index on disk, while the MetaGraph representation expands manyfold to improve query throughput via batch queries. At the same time, Mac-dBG provides query speed as fast as the fastest existing c-dBG index, Fulgor. This enhanced trade-off can be visualized in Fig. 3. We believe these characteristics make Mac-dBG a very promising data structure for enabling large-scale color queries across a range of different applications.

**Fig. 3:**
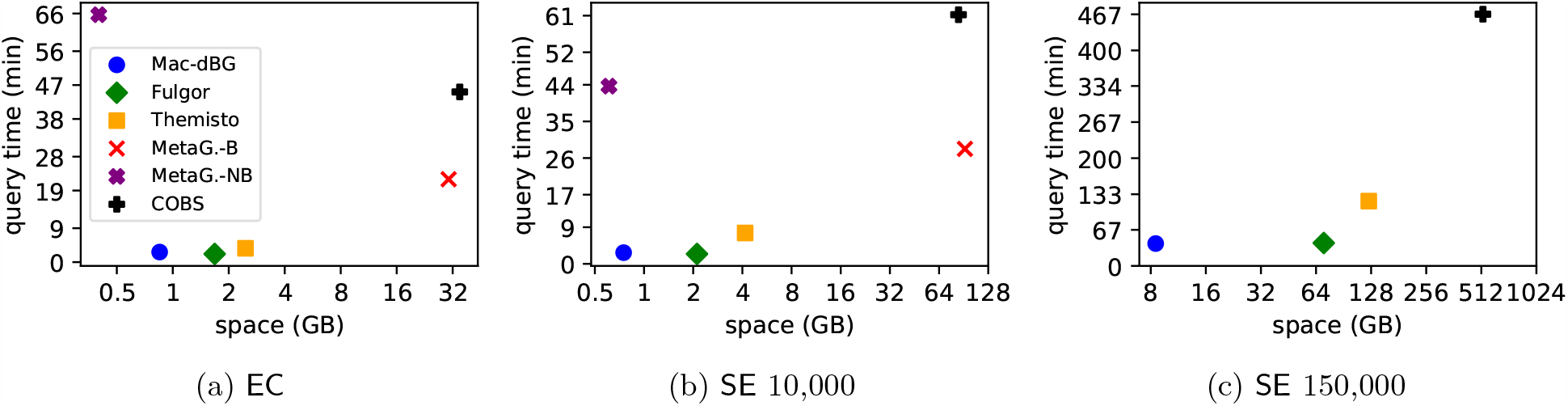
The same data from Table 2 but shown as space vs. time trade-off curves, for some example datasets.

To achieve these substantial improvements over the prior state of the art, the Mac-dBG focuses on providing an improved representation of the color table, the element of the index that tends to grow most quickly as the number of indexed references increases. Specifically, Mac-dBG compresses the colors by factoring out shared sub-patterns that occur across different colors. The color table is represented as a set of *meta* colors and *partial* colors which are combined to recover the original colors exactly. While most interesting formulations of determining the optimal factorization into meta colors and partial colors appear NP-hard, we nonetheless describe a heuristic approach that works well in practice.

Future work would first focus on improving the compression of partial colors even further with the help of a more principled permutation *π* and that of the meta colors with a more succinct encoding method. Lastly, we plan to engineer the construction steps of the SCPO framework. We include in the Supplementary material a list of interesting research questions related to the Mac-dBG data structure.

## Acknowledgements

We are grateful to Laxman Dhulipala for useful comments on an early draft of this paper.

## Declarations

R.P. is a co-founder of Ocean Genomics inc.

## References

1. Bo Liu, Hongzhe Guo, Michael Brudno, and Yadong Wang. deBGA: read alignment with de bruijn graph-based seed and extension. Bioinformatics, 32(21):3224–3232, July 2016.

2. Nicolas L Bray, Harold Pimentel, Páll Melsted, and Lior Pachter. Near-optimal probabilistic rna-seq quantification. Nature biotechnology, 34(5):525–527, 2016.

3. L Schaeffer, H Pimentel, N Bray, P Melsted, and L Pachter. Pseudoalignment for metagenomic read assignment. Bioinformatics, 33(14):2082–2088, 02 2017.

4. Fatemeh Almodaresi, Hirak Sarkar, Avi Srivastava, and Rob Patro. A space and time-efficient index for the compacted colored de Bruijn graph. Bioinformatics, 34(13):i169–i177, 2018.

5. Mark Reppell and John Novembre. Using pseudoalignment and base quality to accurately quantify microbial community composition. PLOS Computational Biology, 14(4):1–23, 04 2018.

6. Tommi Mäklin, Teemu Kallonen, Sophia David, Christine J Boinett, Ben Pascoe, Guillaume Méric, David M Aanensen, Edward J Feil, Stephen Baker, Julian Parkhill, et al. High-resolution sweep metagenomics using fast probabilistic inference [version 1; peer review: 1 approved, 1 approved with reservations]. Wellcome open research, 5(14), 2021.

7. Fatemeh Almodaresi, Mohsen Zakeri, and Rob Patro. PuffAligner: a fast, efficient and accurate aligner based on the pufferfish index. Bioinformatics, 37(22):4048–4055, June 2021.

8. Giorgos Skoufos, Fatemeh Almodaresi, Mohsen Zakeri, Joseph N. Paulson, Rob Patro, Artemis G. Hatzigeorgiou, and Ioannis S. Vlachos. AGAMEMNON: an accurate metaGenomics and MEtatranscriptoMics quaNtificatiON analysis suite. Genome Biology, 23(1), January 2022.

9. Ilia Minkin and Paul Medvedev. Scalable multiple whole-genome alignment and locally collinear block construction with SibeliaZ. Nature Communications, 11(1), December 2020.

10. Ilia Minkin and Paul Medvedev. Scalable pairwise whole-genome homology mapping of long genomes with BubbZ. iScience, 23(6):101224, June 2020.

11. Siavash Sheikhizadeh, M. Eric Schranz, Mehmet Akdel, Dick de Ridder, and Sandra Smit. PanTools: representation, storage and exploration of pan-genomic data. Bioinformatics, 32(17):i487–i493, August 2016.

12. Alan Cleary, Thiruvarangan Ramaraj, Indika Kahanda, Joann Mudge, and Brendan Mumey. Exploring Frequented Regions in Pan-Genomic Graphs. IEEE/ACM Transactions on Computational Biology and Bioinformatics, 16(5):1424–1435, September 2019.

13. Buwani Manuweera, Joann Mudge, Indika Kahanda, Brendan Mumey, Thiruvarangan Ramaraj, and Alan Cleary. Pangenome-Wide Association Studies with Frequented Regions. In Proceedings of the 10th ACM International Conference on Bioinformatics, Computational Biology and Health Informatics. ACM, September 2019.

14. Kadir Dede and Enno Ohlebusch. Dynamic construction of pan-genome subgraphs. Open Computer Science, 10(1):82–96, April 2020.

15. John A. Lees, T. Tien Mai, Marco Galardini, Nicole E. Wheeler, Samuel T. Horsfield, Julian Parkhill, and Jukka Corander. Improved Prediction of Bacterial Genotype-Phenotype Associations Using Interpretable Pangenome-Spanning Regressions. mBio, 11(4), August 2020.

16. Roland Wittler. Alignment- and reference-free phylogenomics with colored de Bruijn graphs. Algorithms for Molecular Biology, 15(1), April 2020.

17. Nina Luhmann, Guillaume Holley, and Mark Achtman. BlastFrost: fast querying of 100, 000s of bacterial genomes in bifrost graphs. Genome Biology, 22(1), January 2021.

18. Amatur Rahman, Yoann Dufresne, and Paul Medvedev. Compression Algorithm for Colored de Bruijn Graphs. In 23rd International Workshop on Algorithms in Bioinformatics (WABI 2023), pages 17:1–17:14, 2023.

19. Shoshana Marcus, Hayan Lee, and Michael C Schatz. Splitmem: a graphical algorithm for pan-genome analysis with suffix skips. Bioinformatics, 30(24):3476–3483, 2014.

20. Uwe Baier, Timo Beller, and Enno Ohlebusch. Graphical pan-genome analysis with compressed suffix trees and the burrows–wheeler transform. Bioinformatics, 32(4):497–504, 2016.

21. Jason Fan, Noor Pratap Singh, Jamshed Khan, Giulio Ermanno Pibiri, and Rob Patro. Fulgor: A Fast and Compact k-mer Index for Large-Scale Matching and Color Queries. In 23rd International Workshop on Algorithms in Bioinformatics (WABI 2023), pages 18:1–18:21, 2023. URL: https://github.com/jermp/fulgor.

22. Justin Zobel and Alistair Moffat. Inverted files for text search engines. ACM Computing Surveys (CSUR), 38(2):6–es, 2006.

23. Giulio Ermanno Pibiri and Rossano Venturini. Techniques for inverted index compression. ACM Computing Surveys (CSUR), 53(6):125:1–125:36, 2021.

24. Jason Fan, Jamshed Khan, Giulio Ermanno Pibiri, and Rob Patro. Spectrum preserving tilings enable sparse and modular reference indexing. In Research in Computational Molecular Biology, pages 21–40, 2023.

25. Giulio Ermanno Pibiri. Sparse and skew hashing of k-mers. Bioinformatics, 38(Supplement 1):i185–i194, 06 2022.

26. Camille Marchet, Christina Boucher, Simon J Puglisi, Paul Medvedev, Mikaël Salson, and Rayan Chikhi. Data structures based on k-mers for querying large collections of sequencing data sets. Genome Research, 31(1):1–12, 2021.

27. Jarno N Alanko, Jaakko Vuohtoniemi, Tommi Mäklin, and Simon J Puglisi. Themisto: a scalable colored k-mer index for sensitive pseudoalignment against hundreds of thousands of bacterial genomes. Bioinformatics, 39(Supplement 1):i260–i269, June 2023. URL: https://github.com/algbio/themisto.

28. Jarno N. Alanko, Simon J. Puglisi, and Jaakko Vuohtoniemi. Small searchable k-spectra via subset rank queries on the spectral burrows-wheeler transform. SIAM Conference on Applied and Computational Discrete Algorithms (ACDA23), pages 225–236, 2023.

29. Mikhail Karasikov, Harun Mustafa, Amir Joudaki, Sara Javadzadeh-no, Gunnar Rätsch, and André Kahles. Sparse Binary Relation Representations for Genome Graph Annotation. Journal of Computational Biology, 27(4):626–639, April 2020. URL: https://github.com/ratschlab/metagraph.

30. Alexander Bowe, Taku Onodera, Kunihiko Sadakane, and Tetsuo Shibuya. Succinct de Bruijn graphs. In International Workshop on Algorithms in Bioinformatics (WABI), pages 225–235. Springer, 2012.

31. Mikhail Karasikov, Harun Mustafa, Gunnar Rätsch, and André Kahles. Lossless indexing with counting de bruijn graphs. Genome Research, 32(9):1754–1764, 2022.

32. Guillaume Holley and Páll Melsted. Bifrost: highly parallel construction and indexing of colored and compacted de Bruijn graphs. Genome biology, 21(1):1–20, 2020.

33. Daniel Lemire, Owen Kaser, Nathan Kurz, Luca Deri, Chris O’Hara, Francois Saint-Jacques, and Gregory Ssi-Yan-Kai. Roaring bitmaps: Implementation of an optimized software library. Software: Practice and Experience, 48(4):867–895, 2018.

34. Timo Bingmann, Phelim Bradley, Florian Gauger, and Zamin Iqbal. Cobs: a compact bit-sliced signature index. In International Symposium on String Processing and Information Retrieval, pages 285–303. Springer, 2019. URL: https://github.com/bingmann/cobs.

35. Giulio Ermanno Pibiri. On weighted k-mer dictionaries. Algorithms for Molecular Biology, 18(3), 2023.

36. Laxman Dhulipala, Igor Kabiljo, Brian Karrer, Giuseppe Ottaviano, Sergey Pupyrev, and Alon Shalita. Compressing graphs and indexes with recursive graph bisection. In Proceedings of the 22nd ACM SIGKDD International Conference on Knowledge Discovery and Data Mining, pages 1535–1544, 2016.

37. Flavio Chierichetti, Ravi Kumar, Silvio Lattanzi, Michael Mitzenmacher, Alessandro Panconesi, and Prabhakar Raghavan. On compressing social networks. In Proceedings of the 15th ACM SIGKDD international conference on Knowledge discovery and data mining, pages 219–228, 2009.

38. L. H. Harper. Optimal assignments of numbers to vertices. Journal of the Society for Industrial and Applied Mathematics, 12(1):131–135, March 1964.

39. Michael R. Garey and David S. Johnson. Computers and Intractability; A Guide to the Theory of NP-Completeness. W. H. Freeman & Co., USA, 1990.

40. Philippe Flajolet, Éric Fusy, Olivier Gandouet, and Frédéric Meunier. Hyperloglog: the analysis of a near-optimal cardinality estimation algorithm. In Discrete Mathematics and Theoretical Computer Science, pages 137–156. Discrete Mathematics and Theoretical Computer Science, 2007.

41. Karel Břinda, Leandro Lima, Simone Pignotti, Natalia Quinones-Olvera, Kamil Salikhov, Rayan Chikhi, Gregory Kucherov, Zamin Iqbal, and Michael Baym. Efficient and Robust Search of Microbial Genomes via Phylogenetic Compression. April 2023.

42. Stephen F Altschul, Warren Gish, Webb Miller, Eugene W Myers, and David J Lipman. Basic local alignment search tool. Journal of Molecular Biology, 215(3):403–410, 1990.

43. Jarno N. Alanko. 3682 E. Coli assemblies from NCBI, 2022. URL: https://zenodo.org/records/6577997.

44. Grace A. Blackwell, Martin Hunt, Kerri M. Malone, Leandro Lima, Gal Horesh, Blaise T. F. Alako, Nicholas R. Thomson, and Zamin Iqbal. Exploring bacterial diversity via a curated and searchable snapshot of archived DNA sequences. PLOS Biology, 19(11):1–16, 11 2021. URL: http://ftp.ebi.ac.uk/pub/databases/ENA2018-bacteria-661k.

45. Pranvera Hiseni, Knut Rudi, Robert C Wilson, Finn Terje Hegge, and Lars Snipen. HumGut: a comprehensive human gut prokaryotic genomes collection filtered by metagenome data. Microbiome, 9(1):1–12, 2021. URL: https://arken.nmbu.no/-larssn/humgut/index.htm.

46. Mikhail Karasikov, Harun Mustafa, Daniel Danciu, Christopher Barber, Marc Zimmermann, Gunnar Rätsch, and André Kahles. Metagraph: Indexing and analysing nucleotide archives at petabase-scale. BioRxiv, pages 2020–10, 2020.

