## Supplementary material. for "Meta-colored compacted de Bruijn graphs"

### Meta-colored compacted de Bruijn graphs — Supplementary material

**Datasets statistics.** Table 4 reports some basic statistics of the tested collections.

**Space breakdowns.** Fig. 4 shows index space breakdowns for Mac-dBG in comparison to Fulgor. We remark that the dictionary component (the SSHash data structure) is common to both Mac-dBG's and Fulgor's implementations. As evident, the fraction of space for the colors is significantly reduced in Mac-dBG compared to Fulgor — sometimes, even dramatically. Consider, for example, the SE 50,000 pangenome: while the colors take 93% of the whole Fulgor index space, this percentage is now reduced to $(48+15)\% = 63\%$ for Mac-dBG, surprisingly making SSHash a "heavy" component of the index. Another important consideration regards how the total space for the colors is divided among partial and meta colors. Meta colors take even more space than partial colors for larger collections, as apparent from SE 50,000, 100,000, and 150,000 — surely a consequence of the simple $\log_2(N_p)$-bit encoding we are currently using. We are confident that more succinct encodings can substantially reduce the space for the meta colors.

**Low-hit workloads.** In Table 5 we report the query efficiency of the indexes on low-hit workloads, using the very same experimental methodology as used for the experiment in Table 2. We use the reads from the accession SRR896663 in FASTQ format to simulate a low-hit workload.

As apparent from the timings reported, low-hit workloads can be processed much more quickly than high-hit ones because many $k$-mers are not even found in the index dictionary and intersections are performed for a handful of lists. Nonetheless, the trend already discussed in Section 5 does not change: Mac-dBG is as fast as or even faster than Fulgor. Themisto is slower then Fulgor but competitive, whereas both MetaGraph and COBS perform substantially worse.

**Using a general-purpose compressor atop Mac-dBG.** Table 6 shows the effect of compressing the Mac-dBG with the general-purpose Zstd compressor [1]. The purpose of the experiment is two-fold: (1) obtain a smaller on-disk representation to compare against the original Mac-dBG and the smallest evaluated index, MetaGraph; (2) quantify the compression gains achievable by a Lempel-Ziv-based approach whose compression ratio critically depends on the repetitiveness of the input data.

We can see that the use of Zstd atop Mac-dBG offers an overall good reduction in space. Obviously, the reduction is larger on larger indexes. This suggests that the Mac-dBG still presents some redundancy — this is consistent with the observation that both meta color lists and partial colors within a partition are similar. This similarity is exploited by Zstd but not yet by the encodings that we use in the Mac-dBG.

The net result of the experiment is that using Zstd essentially bridges the gap between the size on-disk of MetaGraph with that of the Mac-dBG for the SE pangenomes with a negligible decompression overhead that would be paid prior to answering queries. Instead, the gap in space is still significant on the EC and GB collections. Future work will thus focus on closing this gap without sacrificing the fast query time of Mac-dBG.

**Open questions.** We include here a list of research questions (both theoretical and practical) about the introduced Mac-dBG representation that are likely worthy of investigation in the near future.

RQ 1. *Do different sketching/clustering algorithms have a meaningful impact on the index size?*
In our current results from Section 5 we use a divisive $K$-means clustering algorithm over hyper-log-log sketches, but other clustering/sketching algorithm combinations could be tested, and may yield meaningfully different solutions to the optimization problem.

Table 4: Basic statistics for the tested collections.

| | *E. Coli* (EC) | *S. Enterica* (SE) | | | | | Gut bacteria (GB) |
|---|---|---|---|---|---|---|---|
| Genomes | 3,682 | 5,000 | 10,000 | 50,000 | 100,000 | 150,000 | 30,691 |
| Distinct colors ($\times 10^6$) | 5.59 | 2.69 | 4.24 | 13.92 | 19.36 | 23.61 | 227.80 |
| Integers in colors ($\times 10^9$) | 5.74 | 5.77 | 15.68 | 133.49 | 303.53 | 490.04 | 10.04 |
| $k$-mers in dBG ($\times 10^6$) | 170.65 | 104.69 | 239.88 | 806.23 | 1,018.69 | 1,194.44 | 13,936.86 |
| Unitigs in dBG ($\times 10^6$) | 9.31 | 4.95 | 8.24 | 30.64 | 41.16 | 49.60 | 566.39 |

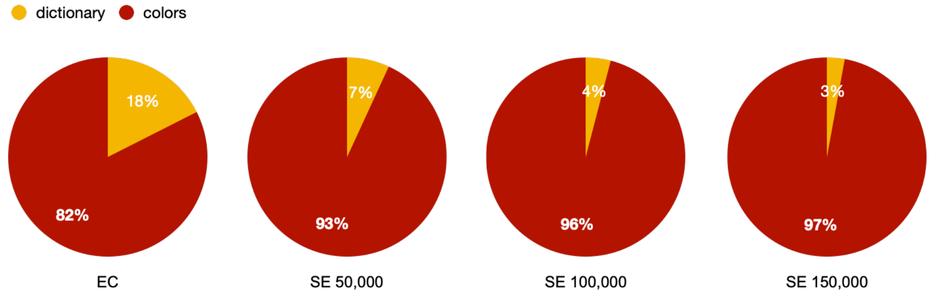


**(a)** Fulgor

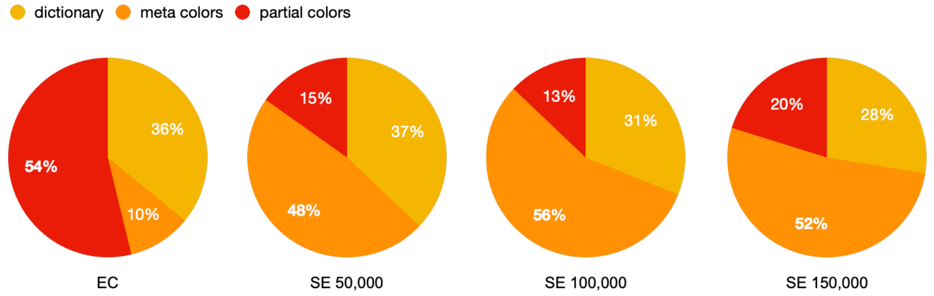


**(b)** Mac-dBG

Fig. 4: Index space breakdowns.

RQ 2. *How should reference identifiers be assigned to references within the same cluster $\mathcal{R}_i$?*
In our current results, we do not impose any particular order on the references within the same cluster; we just follow the input order. However, one can likely better-optimize these assignments to reduce the index size adopting, e.g., techniques similar to optimal leaf re-ordering [2] or recursive bipartite graph partitioning [3] based on the reference similarities already computed from the sketches used for partitioning.

RQ 3. *Can index space be reduced even more by exploiting the fact that partial colors (i.e., colors within the same partition) are similar?*
There is strong evidence from prior work [4,5] that coherence or similarity *between* distinct colors allows for effective compression in a manner that is different from what is obtained via the current SCPO framework. A natural question is the degree to which such approaches can be effectively "stacked", so that one could apply the SCPO framework, and then, within each partition, further compress the partial colors using a referential encoding like the minimum spanning tree-based encoding of Almodaresi et al. [4] or the delta encoding of Karasikov et al. [5]. Unlike the current meta-color scheme, these approaches can reduce the query speed, and so the potential space-time trade-offs are worth exploring. Further, one should understand whether it is more convenient to: (1) compress the partial colors with an entirely separate method, or (2) use the Mac-dBG representation *recursively* on each color partition $\mathcal{P}_i$.

Table 5: Query results for low-hit workloads. We use the same experimental methodology for this experiment as for the one described in the caption of Table 2.

| | Genomes | Rate | Mac-dBG | | Fulgor | | Themisto | | MetaG.-B | | MetaG.-NB | | COBS | |
|---|---|---|---|---|---|---|---|---|---|---|---|---|---|---|
| | | | m:ss | GB | m:ss | GB | m:ss | GB | m:ss | GB | m:ss | GB | mm:ss | GB |
| EC | 3,682 | 4.71 | 0:09 | 0.83 | 0:10 | 1.65 | 0:33 | 2.35 | 7:34 | 2.82 | 3:40 | 0.38 | 10:25 | 28.94 |
| SE | 5,000 | 1.27 | 0:09 | 0.35 | 0:09 | 0.77 | 0:30 | 1.77 | 6:48 | 2.76 | 2:55 | 0.31 | 11:50 | 37.64 |
| | 10,000 | 13.86 | 0:09 | 0.70 | 0:10 | 2.01 | 0:36 | 4.06 | 7:35 | 3.00 | 4:17 | 0.56 | 14:33 | 75.63 |
| | 50,000 | 32.64 | 0:12 | 3.32 | 0:25 | 17.91 | 0:56 | 33.07 | 8:33 | 5.05 | 6:47 | 2.42 | 39:33 | 367.34 |
| | 100,000 | 34.09 | 0:14 | 5.60 | 0:45 | 41.49 | 1:22 | 75.89 | 9:19 | 7.04 | 7:33 | 4.23 | 48:52 | 521.58 |
| | 150,000 | 34.06 | 0:16 | 7.41 | 1:06 | 69.05 | 2:05 | 124.19 | — | — | — | — | 37:40 | 522.47 |
| GB | 30,691 | 11.77 | 0:46 | 28.51 | 0:57 | 36.02 | 1:42 | 48.37 | 11:03 | 12.34 | 11:55 | 9.89 | 30:01 | 192.70 |

Table 6: Zstd compression algorithm applied to the Mac-dBG along with decompression time, in comparison to the space of Mac-dBG and MetaGraph.

| | Genomes | Mac-dBG | Mac-dBG + Zstd | | MetaGraph |
|---|---|---|---|---|---|
| | | Total GB | Total GB | decompr. (sec) | Total GB |
| EC | 3,682 | 0.81 | 0.73 (−9.88%) | 0.7 | 0.33 |
| SE | 5,000 | 0.32 | 0.27 (−15.63%) | 0.4 | 0.26 |
| | 10,000 | 0.68 | 0.59 (−13.24%) | 0.7 | 0.51 |
| | 50,000 | 3.40 | 2.09 (−38.53%) | 3.2 | 2.31 |
| | 100,000 | 5.55 | 4.25 (−23.42%) | 4.9 | 3.95 |
| | 150,000 | 7.40 | 5.62 (−24.05%) | 7.0 | — |
| GB | 30,691 | 29.16 | 26.99 (−7.44%) | 29.00 | 10.00 |

RQ 4. *How do different compression methods impact the space/time trade-offs of* Mac-dBG*s?*
Clearly, there is no single compression algorithm that works best for all needs; rather, one should choose a space/time trade-off suitable for the application at hand. In principle, one could completely ignore query efficiency and just support decoding of the colors in order (i.e., first $C_1$, then $C_2$, and so on) with the purpose of optimizing index space. This extreme trade-off — sometimes called "disk" compression — is gaining popularity in recent years given the massive size of genomic collections. The goal of such algorithms [6,7] is therefore to save disk space/energy and the sharing of large datasets more economical.

RQ 5. *What is the information-theoretic space lower bound on the representation of any* Mac-dBG*?*
This is a question of great importance. On one hand, lower bounds exist for the (arguably simpler) case of un-colored de Bruijn graphs, both for more general membership data structures [8] and navigational data structures [9]. Moreover, practical data structures exist to represent de Bruijn graphs using only a few bits per $k$-mer and supporting membership [10,11,12] even coming close to the navigational space lower bound [12]. On the other hand, there is a distinct lack of similar information theoretic lower bounds on the representation of the c-dBG. Yet, this appears to be a needed tool to understand how much improvement might be expected, and what size of representation might be achieved.

#### References

1. Meta Platforms Inc. Zstandard: Fast real-time compression algorithm, 2023. URL: https://github.com/facebook/zstd/releases/tag/v1.5.5.
2. Ziv Bar-Joseph, David K. Gifford, and Tommi S. Jaakkola. Fast optimal leaf ordering for hierarchical clustering. *Bioinformatics*, 17(suppl_1):S22–S29, June 2001.
3. Laxman Dhulipala, Igor Kabiljo, Brian Karrer, Giuseppe Ottaviano, Sergey Pupyrev, and Alon Shalita. Compressing graphs and indexes with recursive graph bisection. In *Proceedings of the 22nd ACM SIGKDD International Conference on Knowledge Discovery and Data Mining*, pages 1535–1544, 2016.
4. Fatemeh Almodaresi, Prashant Pandey, Michael Ferdman, Rob Johnson, and Rob Patro. An Efficient, Scalable, and Exact Representation of High-Dimensional Color Information Enabled Using de Bruijn Graph Search. *Journal of Computational Biology*, 27(4):485–499, April 2020.
5. Mikhail Karasikov, Harun Mustafa, Gunnar Rätsch, and André Kahles. Lossless indexing with counting de bruijn graphs. *Genome Research*, 32(9):1754–1764, 2022.
6. Amatur Rahman, Rayan Chikhi, and Paul Medvedev. Disk compression of k-mer sets. *Algorithms for Molecular Biology*, 16(1):10, 2021.
7. Amatur Rahman, Yoann Dufresne, and Paul Medvedev. Compression Algorithm for Colored de Bruijn Graphs. In *23rd International Workshop on Algorithms in Bioinformatics (WABI 2023)*, pages 17:1–17:14, 2023.
8. Thomas C. Conway and Andrew J. Bromage. Succinct data structures for assembling large genomes. *Bioinformatics*, 27(4):479–486, January 2011.
9. Rayan Chikhi, Antoine Limasset, Shaun Jackman, Jared T. Simpson, and Paul Medvedev. On the Representation of De Bruijn Graphs. *Journal of Computational Biology*, 22(5):336–352, May 2015.
10. Alexander Bowe, Taku Onodera, Kunihiko Sadakane, and Tetsuo Shibuya. Succinct de Bruijn graphs. In *International Workshop on Algorithms in Bioinformatics (WABI)*, pages 225–235. Springer, 2012.
11. Giulio Ermanno Pibiri. Sparse and skew hashing of k-mers. *Bioinformatics*, 38(Supplement_1):i185–i194, 06 2022.
12. Jarno N. Alanko, Simon J. Puglisi, and Jaakko Vuohtoniemi. Small searchable k-spectra via subset rank queries on the spectral burrows-wheeler transform. *SIAM Conference on Applied and Computational Discrete Algorithms (ACDA23)*, pages 225–236, 2023.
